## Supplementary material for "Identification of transporters essential for survival of *Leishmania* promastigotes in the digestive tract of sand flies": All_Supplementary_Files: Supplementary Figure 2.pdf

#### (A) Control PCR amplification of the blasticidin S acetyltransferase (Bla) encoding gene from mutant gDNA

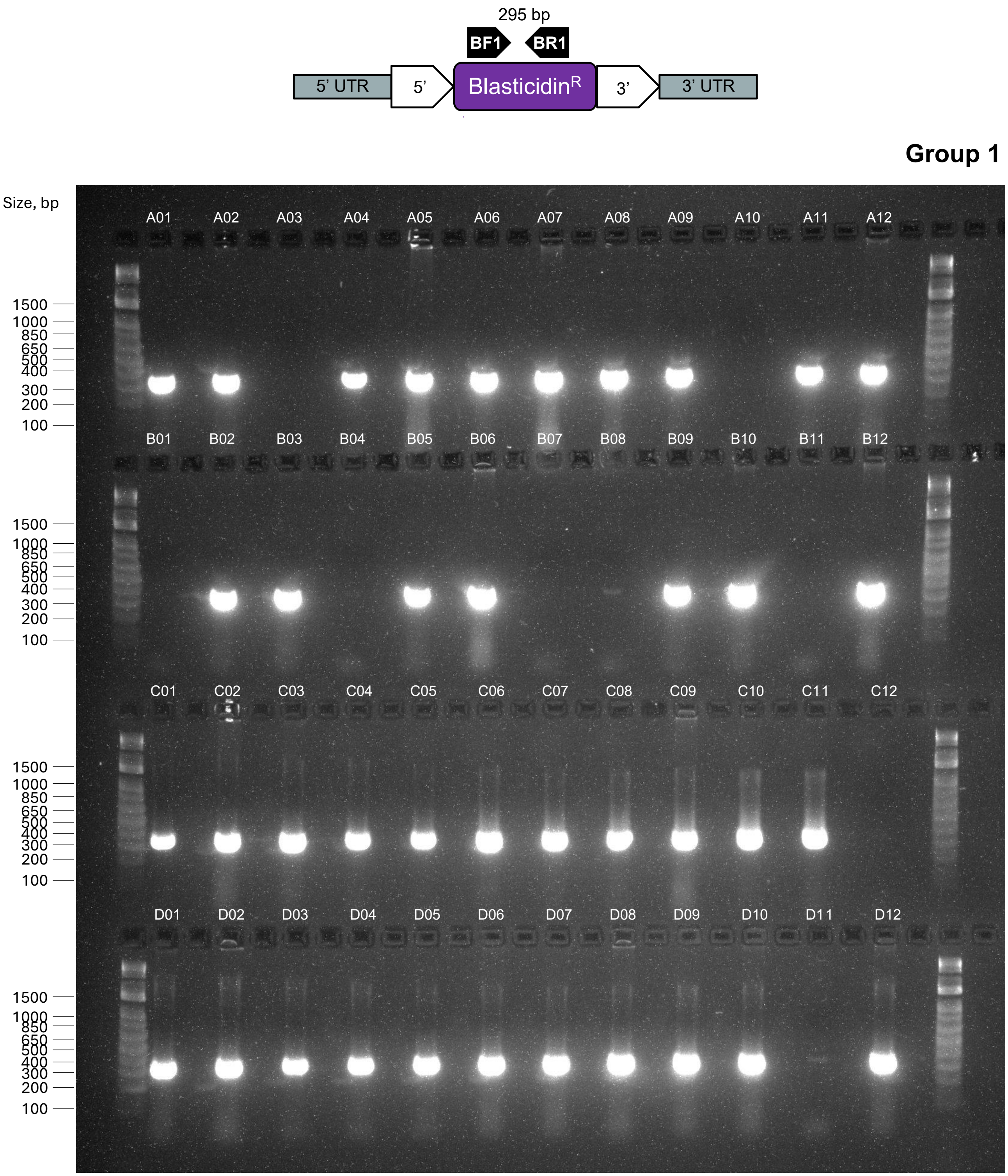


**DNA ladder**
1Kb Plus Dna Ladder, Ref. 10787018, ThermoFisher

### Supplementary Figure 2

#### Group 1

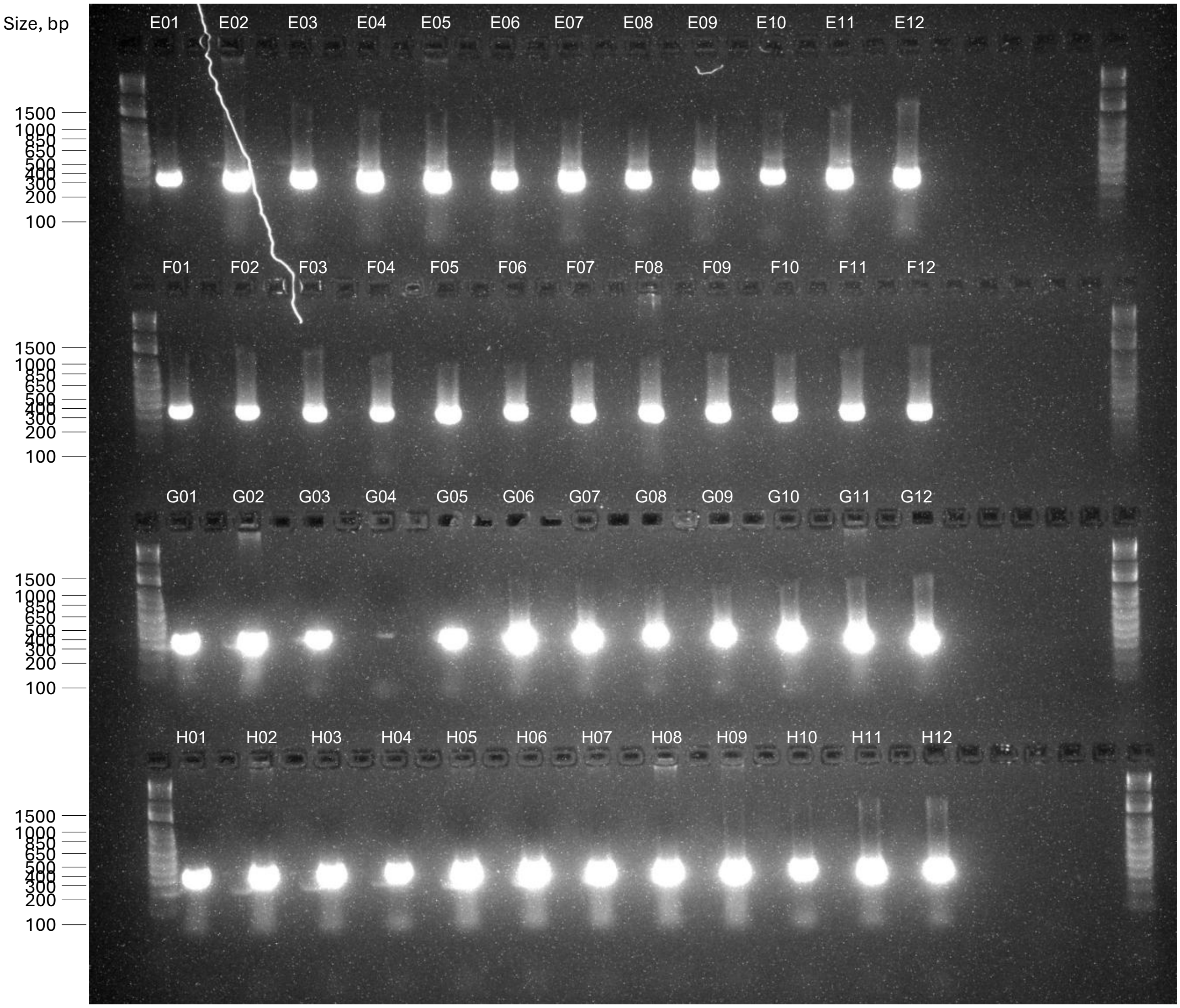


**DNA ladder**

1Kb Plus Dna Ladder, Ref. 10787018, ThermoFisher

#### Group 2

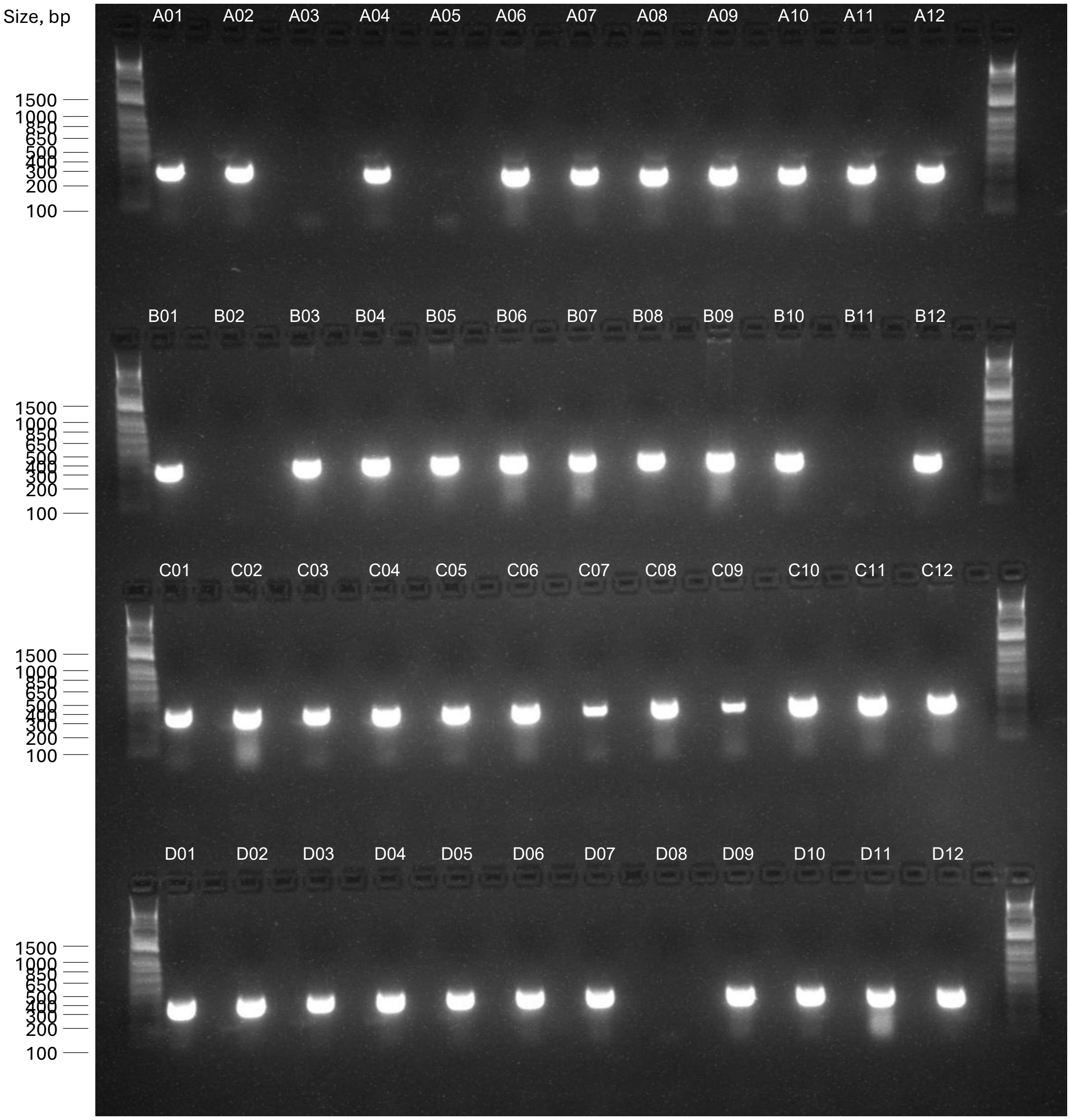


**DNA ladder**

1Kb Plus Dna Ladder, Ref. 10787018, ThermoFisher

#### Group 2

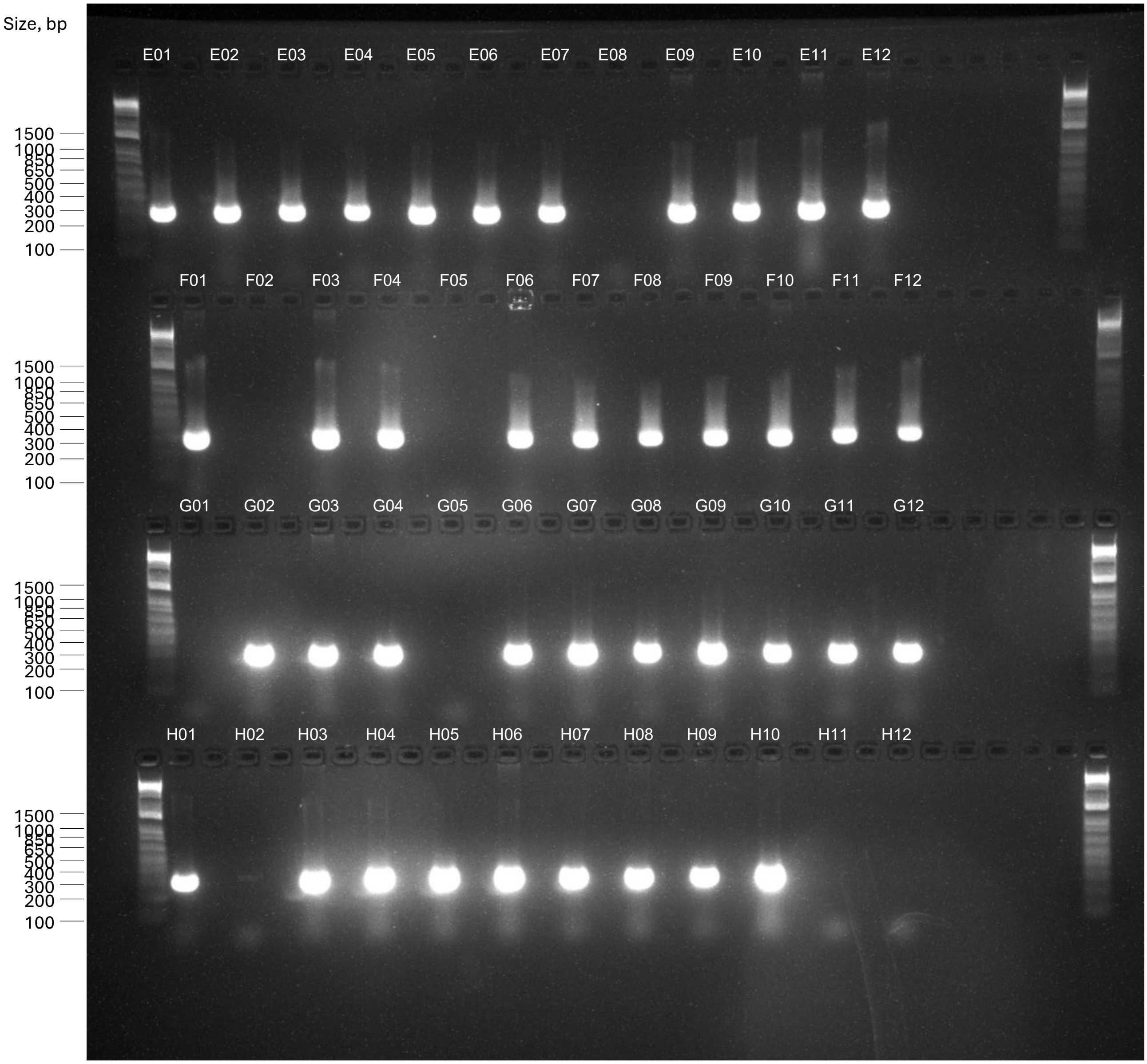


**DNA ladder**

1Kb Plus Dna Ladder, Ref. 10787018, ThermoFisher

#### Group 3

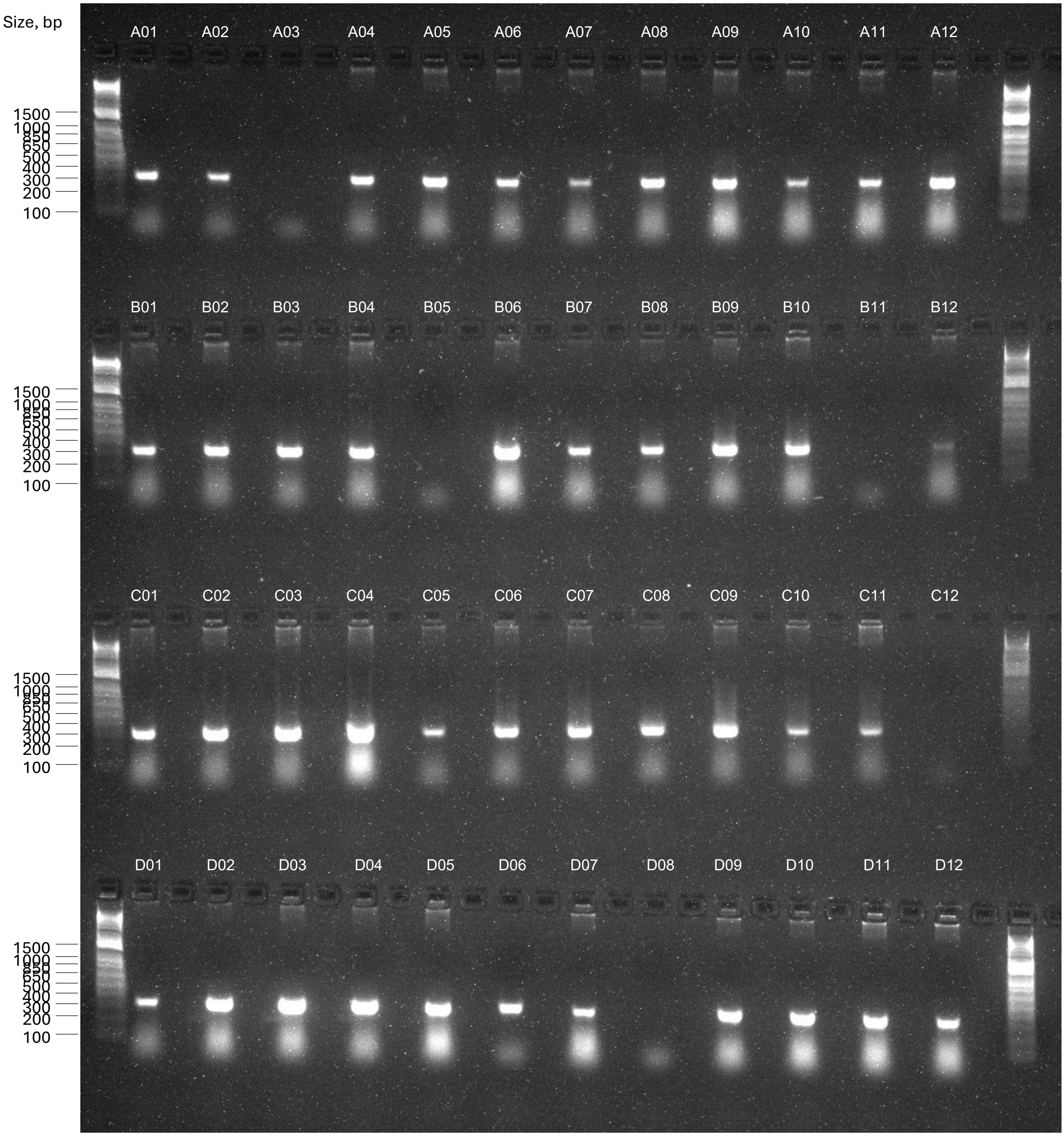


**DNA ladder**

1Kb Plus Dna Ladder, Ref. 10787018, ThermoFisher

#### Group 3

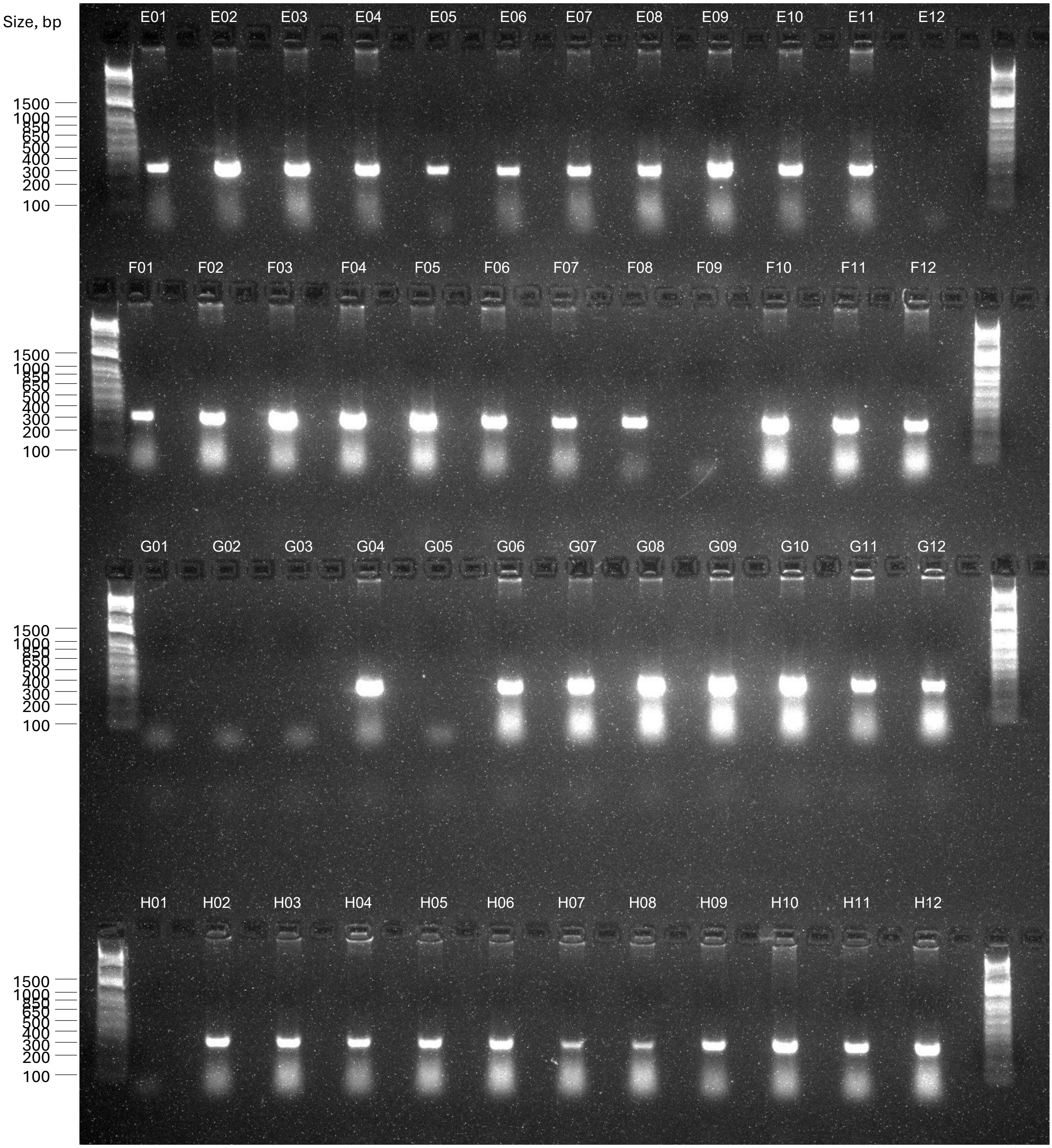


**DNA ladder**

1Kb Plus Dna Ladder, Ref. 10787018, ThermoFisher

#### Group 4

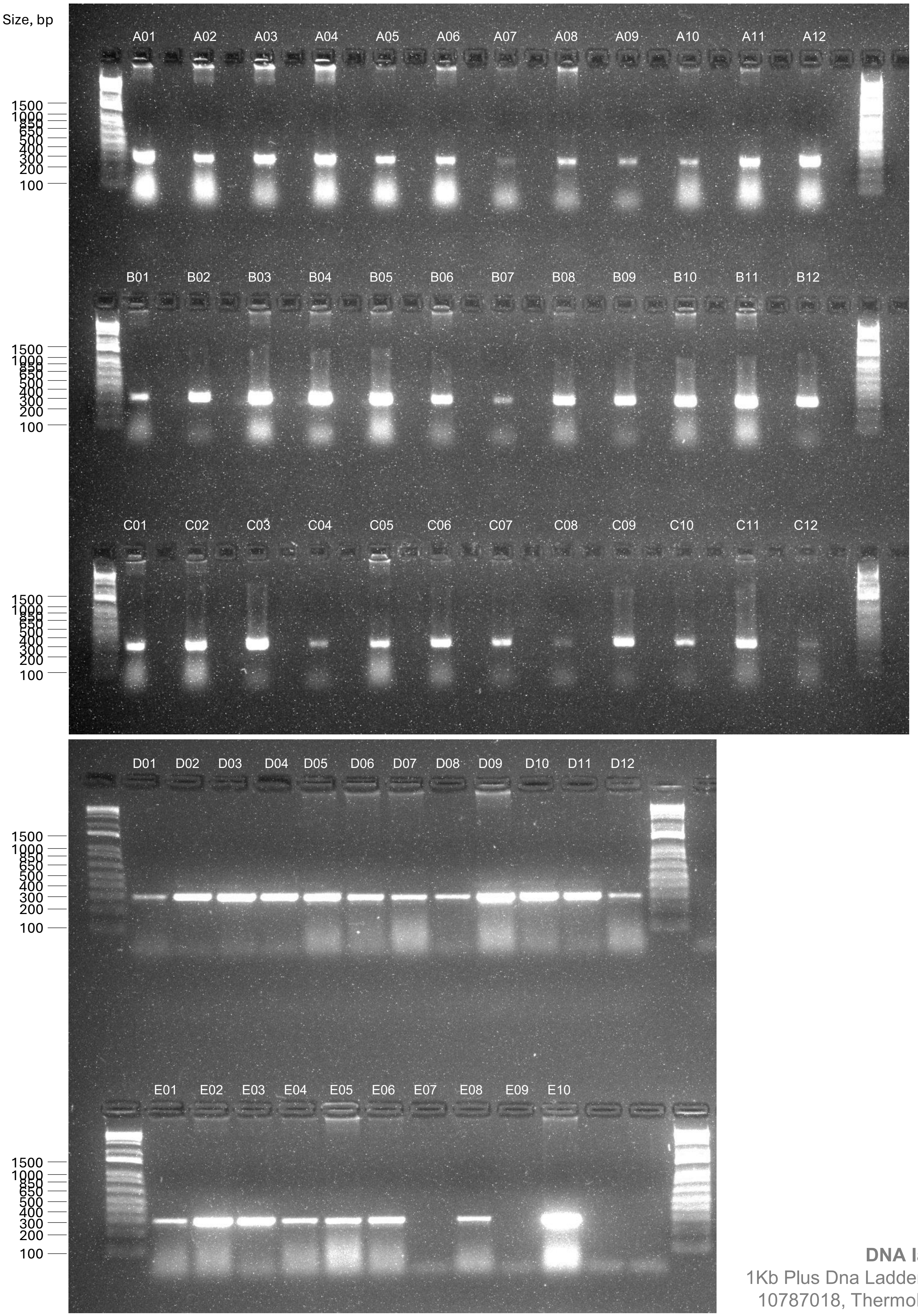


**DNA ladder**
1Kb Plus Dna Ladder, Ref. 10787018, ThermoFisher

#### (B) Diagnostic PCR amplification of target gene for KO validation

**Left lane**, Mutant gDNA
**Right lane**, Parental gDNA (control)

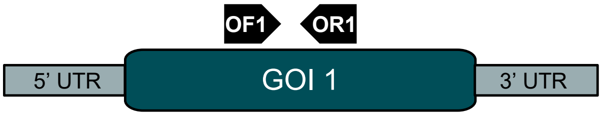


**Group 1**

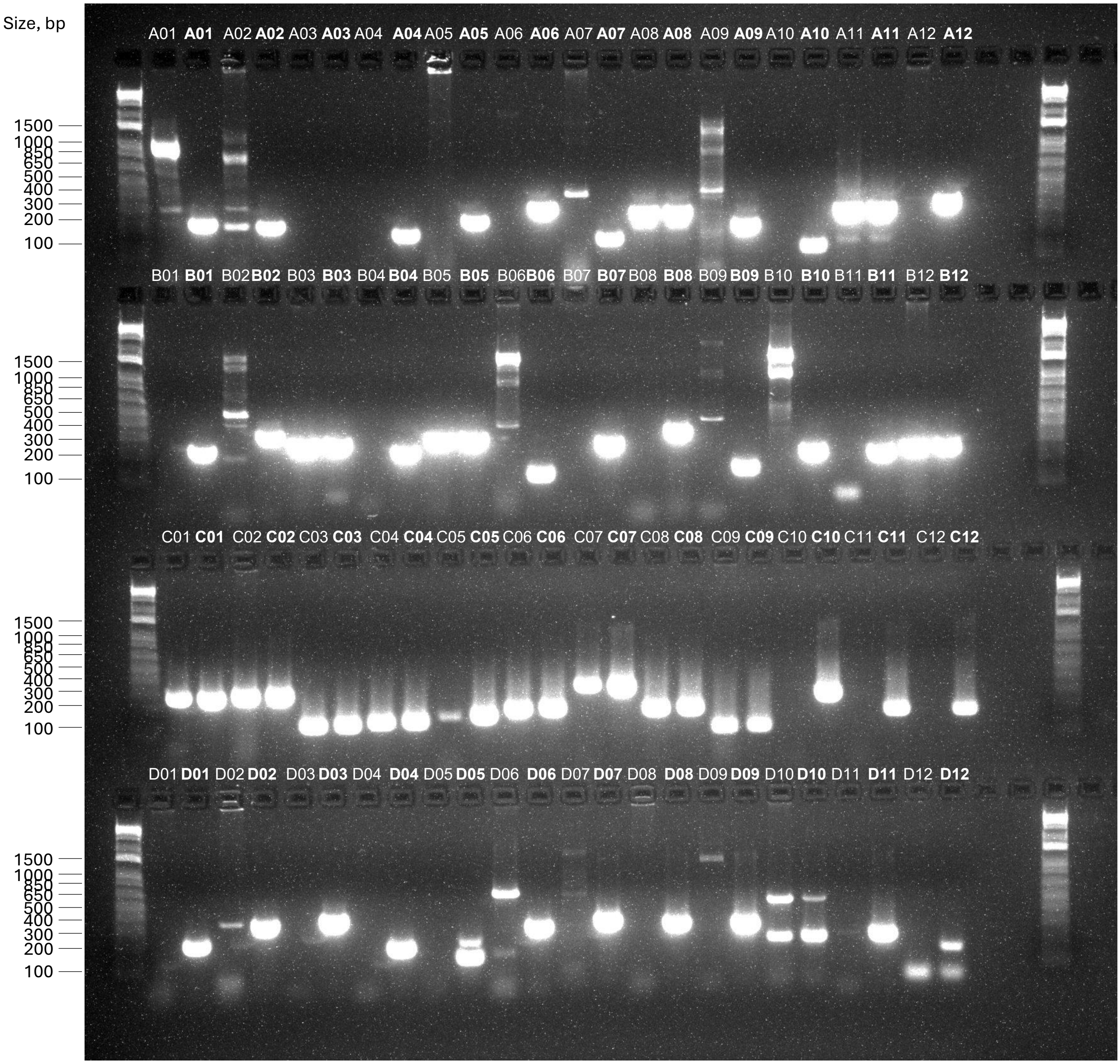


**DNA ladder**
1Kb Plus Dna Ladder, Ref. 10787018, ThermoFisher

#### Group 1

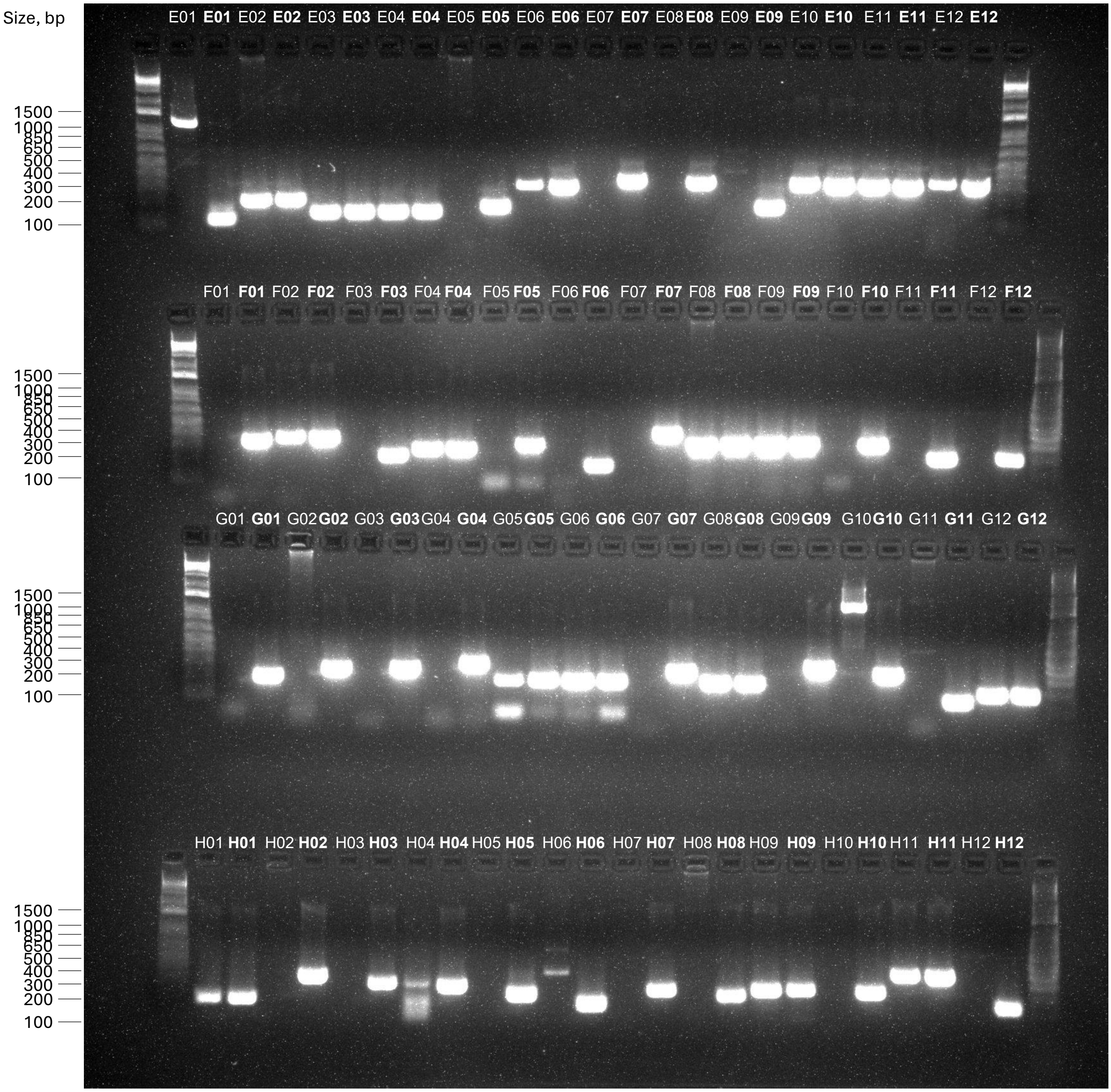


**DNA ladder**

1Kb Plus Dna Ladder, Ref. 10787018, ThermoFisher

#### Group 2

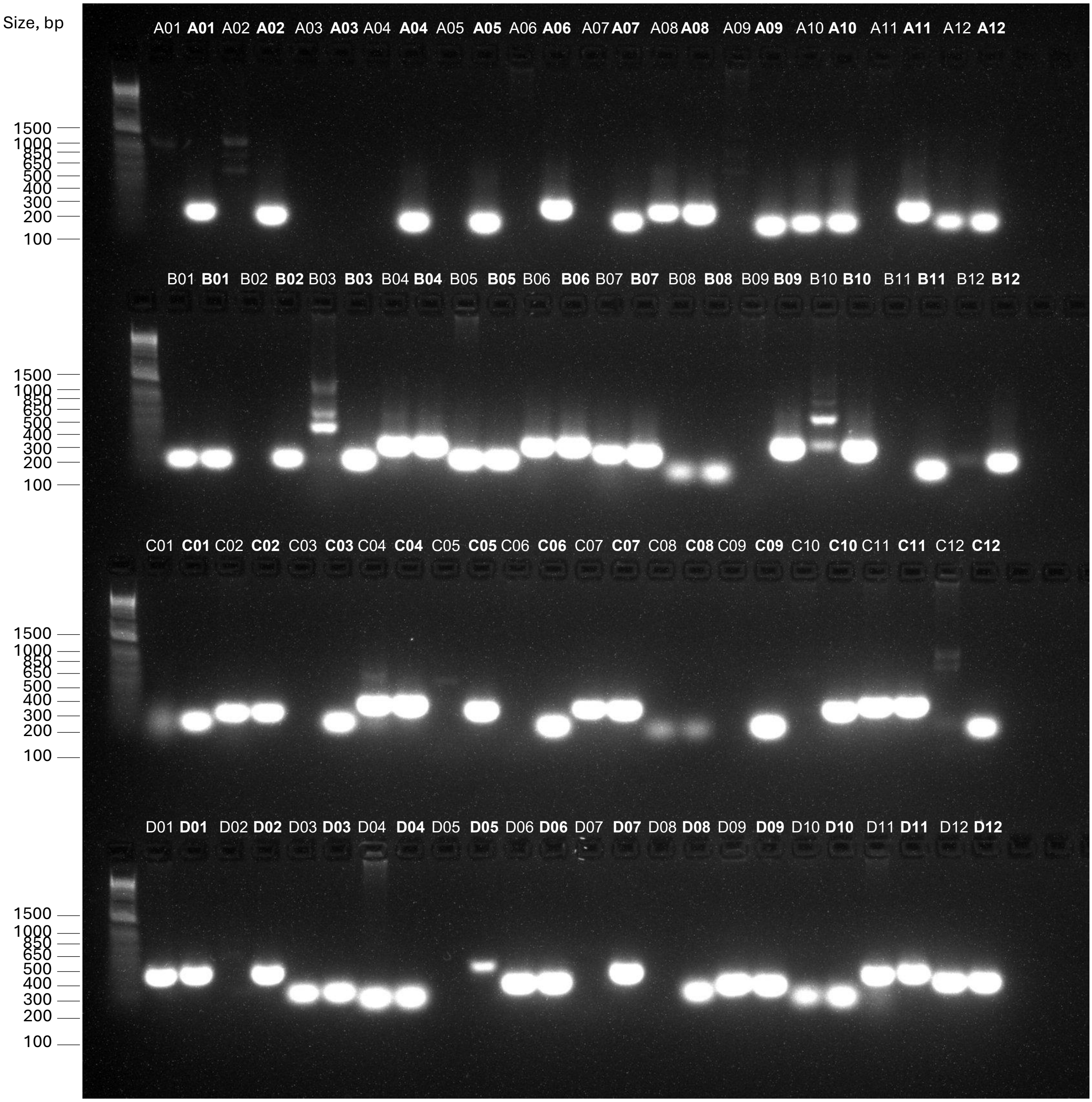


**DNA ladder**

1Kb Plus Dna Ladder, Ref. 10787018, ThermoFisher

#### Group 2

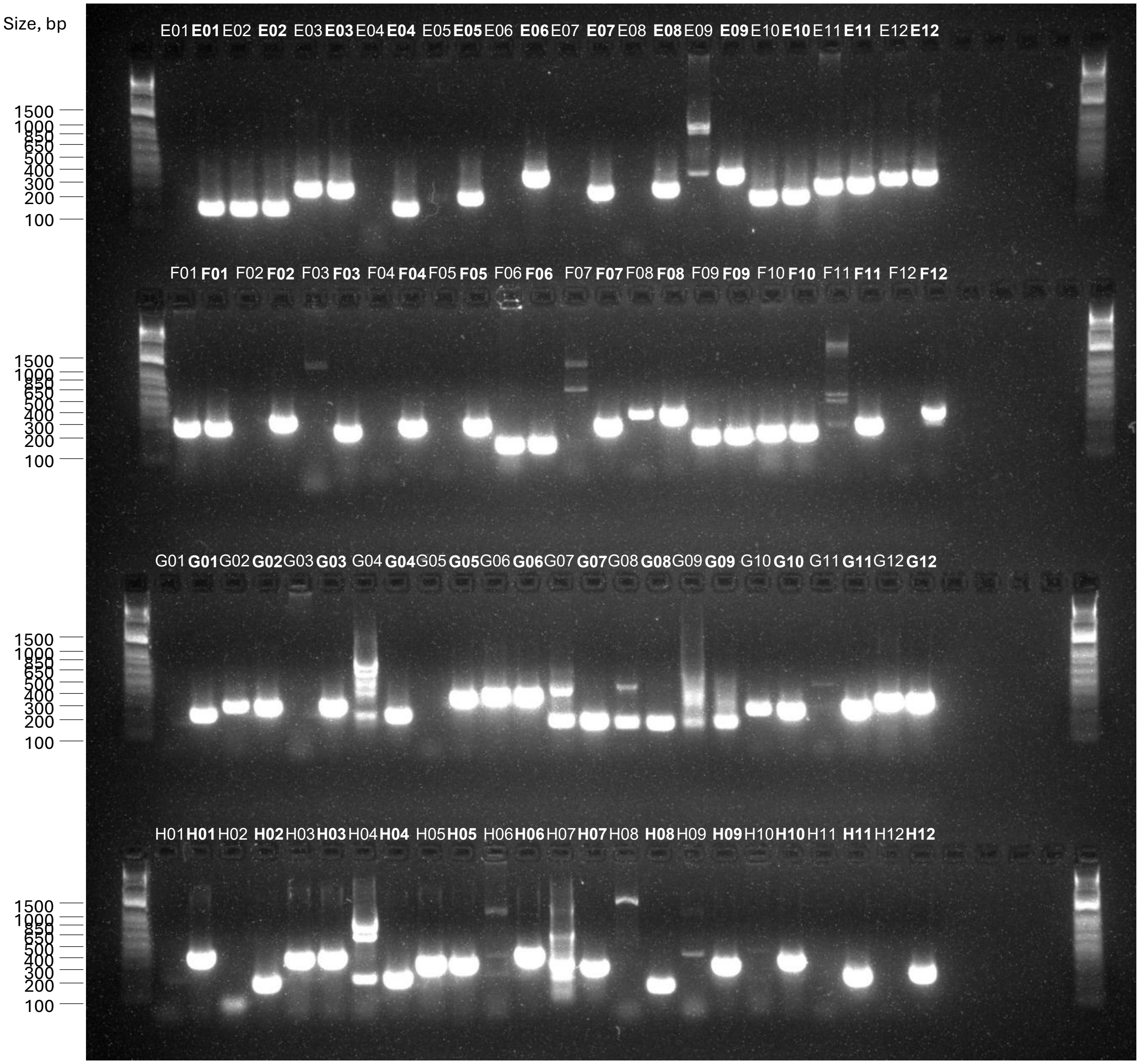


**DNA ladder**
1Kb Plus Dna Ladder, Ref. 10787018, ThermoFisher

#### Group 3

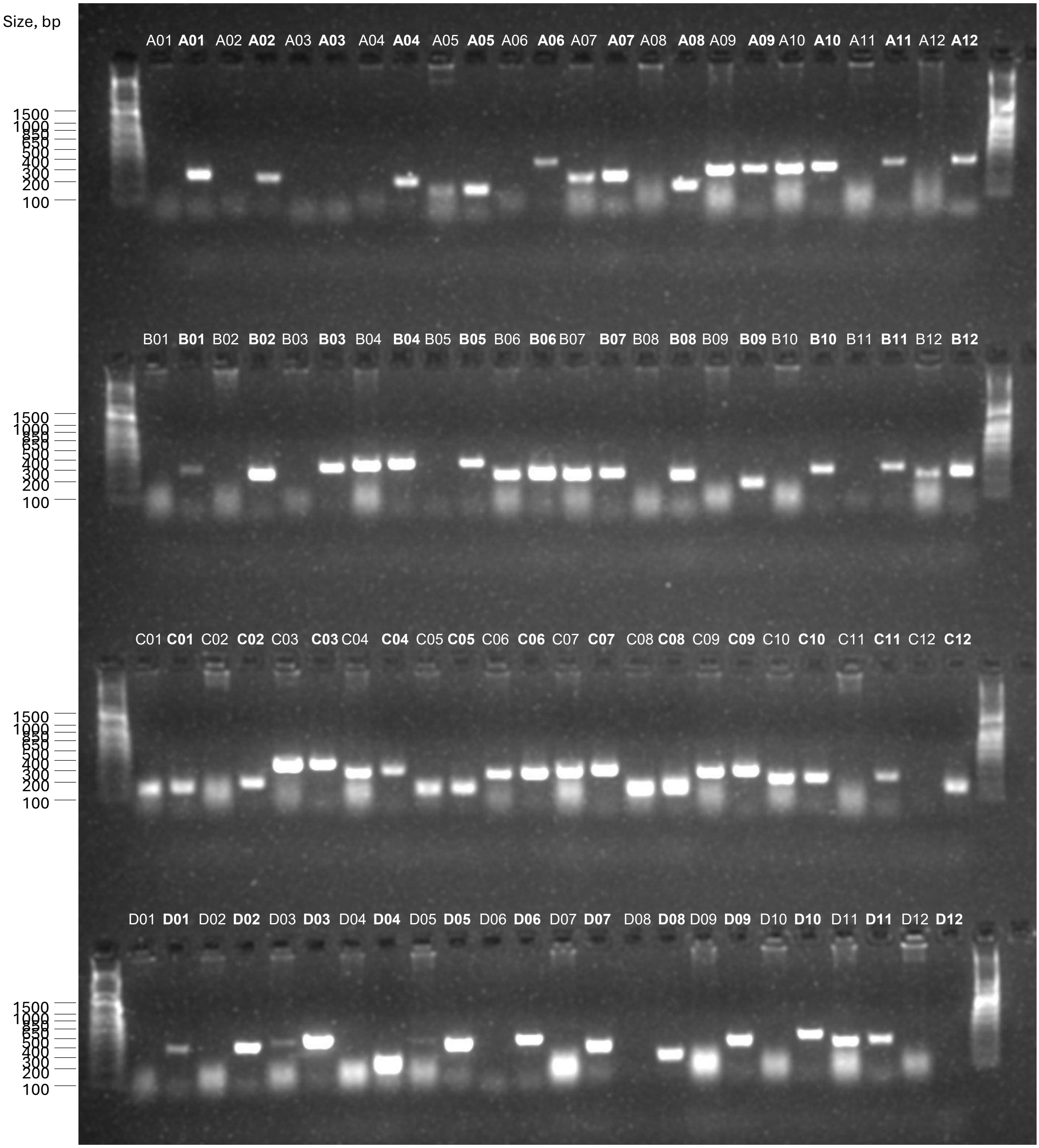


**DNA ladder**

1Kb Plus Dna Ladder, Ref. 10787018, ThermoFisher

#### Group 3

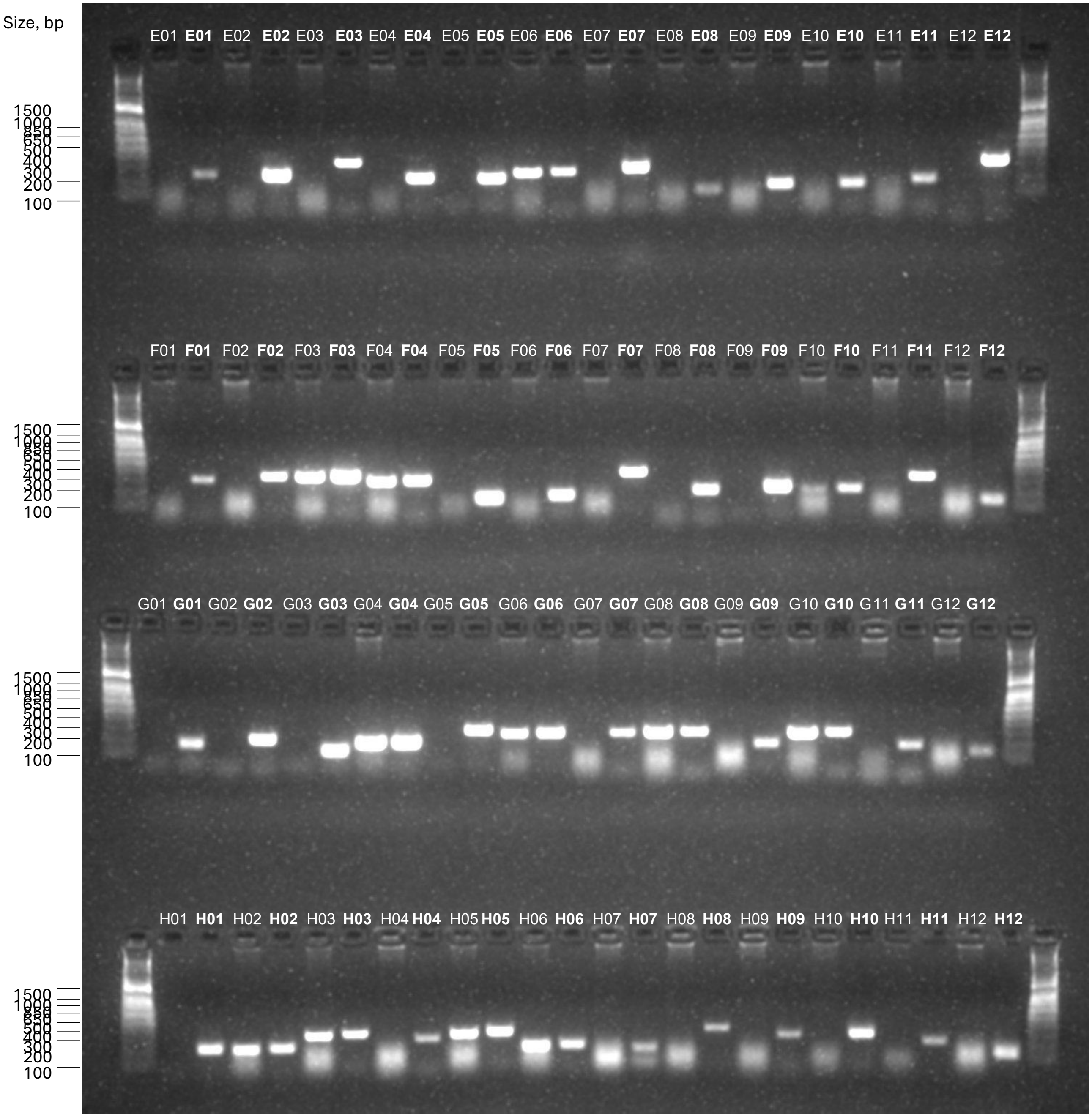


**DNA ladder**
1Kb Plus Dna Ladder, Ref. 10787018, ThermoFisher

#### Group 4

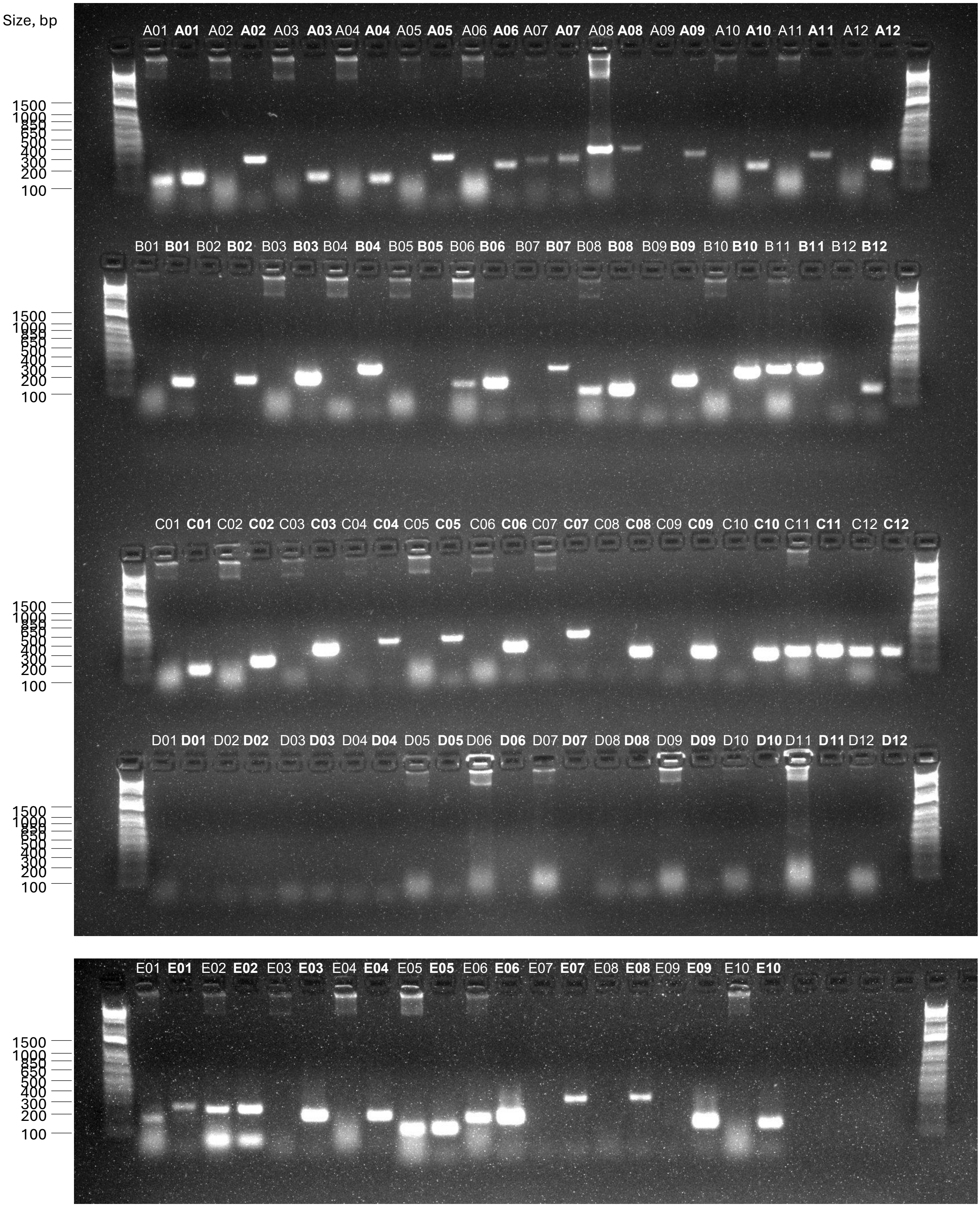


**DNA ladder**

1Kb Plus Dna Ladder, Ref. 10787018, ThermoFisher
