## Supplementary material for "Identification of transporters essential for survival of *Leishmania* promastigotes in the digestive tract of sand flies": All_Supplementary_Files: Supplementary_file_legends.docx

### Description of Supplementary Data

**Supplementary Figure 1. Consolidated summary of gene deletion results for the transportome of *Leishmania mexicana*.**

(A) Top, pie-charts showing the numbers of successful gene deletions (cyan) and non-successful deletion attempts (magenta), across two independent screens (44 and this study). Bottom, break-down of non-successful deletions attempts into two sub-categories: (i) Double drug-resistant populations where ORF is still detected (or PCR inconclusive) (yellow); (ii) Attempts where no drug resistant populations were ever recovered, or populations where resistant cells could only be recovered with single drug selection and ORF was still detected (dark pink). (B) Summary of gene deletion results separated into TCDB families (Supplementary Table 3); colours as for A.

#### Supplementary Figure 2. Diagnostic PCR gel electrophoresis.

Results of diagnostic PCRs for genotypic validation of all new mutant cell lines reported in this study.

#### Supplementary Figure 3. Overview of targeted genes organised in tandem arrays.

(A) Pie-charts summarising gene deletion results separated into TCDB families (left) and in total (right) (Supplementary Table 3). (B) Overview of all genotypes for single genes and whole targeted arrays, coloured in four main categories (as in Supplementary Figure 1).

#### Supplementary Figure 4. Trajectories of the average of normalised reads *in vitro* and *in vivo*.

(A) Plot legends. (B-F) Trajectories of the average of reads for all mutant barcodes in all conditions studied in this report, normalized to total reads for the experiment, reflecting the range of relative barcode proportions for the different mutants in each pool.

#### Supplementary Figure 5. Trajectories of the average of normalised reads *in vitro* and *in vivo*, relative to time zero.

(A) Plot legends. (B-F) Trajectories of the average of barcode reads for all mutants in all conditions studied in this report, normalized to total reads for the experiment and relative to time-point “0 hours” (T_0_).

#### Supplementary Figure 6. Generation, validation and characterisation of the Glucose 1-3 null array mutant.

(A) Diagnostic PCR gel electrophoresis of GT gene array null mutant clones. Lane 1, clone B4 gDNA + GT_array_ORF primers; lane 2, clone B7 gDNA and GT_array_ORF primers; lane 3, clone B4 gDNA and PFR2 primers (positive control for gDNA); lane 4, clone B7 gDNA and PFR2 primers; lane 5, parental gDNA and GT_array_ORF primers; lane 6, parental gDNA and PFR2 primers; lanes 7-9, gDNA of clones B4, B7 and parental without addition of primers. (B) Lane 1, clone H1 and GT_array_ORF primers; lane 2, clone H1 and PFR2 primers; lane 3, parental gDNA and GT_array_ORF primers; lane 4, parental gDNA and PFR2 primers; lanes 5 and 6, negative controls (no gDNA). DNA ladder: GeneRuler 100 bp DNA ladder by Thermo Scientific (REF: SM0243). (C) Locations of primer binding sites for diagnostic PCR. The forward primer (GT_array_ORF_FW) is highlighted with grey circles and the reverse primer (GT_array_ORF_RV) in blue circles. Note that both FW and RV primers recognise all glucose transporter open reading frames (ORFs) in the array. (D) Growth profile of GT KO clones over 72 hours. Black, Parental cell line; Yellow, LmGT array KO clone B4; Turquoise, LmGT array KO clone B7. Data points show the average of three measurements. (E) Doubling times of cells, calculated from D.

#### Supplementary Figure 7. Continuous growth curve of *Leishmania* cell lines.

Growth of parental (PAR, dark blue), V-ATPase V1E null mutant (KO, magenta) and V-ATPase V1E add-back (AB, grey) mutant promastigotes *in vitro* for 5 days of continuous growth. At each time point samples were collected for qPCR analysis (*see Fig 4H*).

**Supplementary File 1. Map of the pTadd-LmxM.36.3100 after restriction digestion cloning.**

FASTA DNA sequence of pTadd-LmxM.36.3100 plasmid used to restore LmxM.36.3100 activity in the ΔLmxM.36.3100 deletion mutant.

#### Supplementary Table 1. – Revised TransLeish DB.

Information about single and array genes targeted in this study.

#### Supplementary Table 2. Barcodes and Primers.

Barcodes and primer sequences used for the generation of mutant cell lines and diagnostic PCRs.

#### Supplementary Table 3. Diagnostic PCR Results.

Results of diagnostic PCRs for knockout validations for all mutant cell lines as well as gel electrophoresis schemes (from Supplementary Figure 1).

#### Supplementary Table 4. Pool membership.

List of barcoded mutant cell lines, barcode sequences and ids included in each analysed pool *in vitro* and *in vivo*.

#### Supplementary Table 5. Fitness Scores and raw read counts.

Fitness scores, p-values from t-test, raw read counts and summary of reads for each timepoint from promastigotes grown *in vitro* (0, 24, 48 and 144 hours) and sand fly infections (0, 48 and 216 hours).

#### Supplementary Table 6. Growth curve raw data and doubling time calculations

Data from densities recorded for the promastigote in vitro cultures, corresponding doubling times, as well as doubling times (calculated from 12).

#### Supplementary Table 7. Cell size measurements from promastigotes in vivo

Measurements of body length, width and flagellar length of Parental, V-ATPase V_1_E null and V-ATPase V_1_E add-back promastigote mutant cell lines isolated at 9 days PBM.

#### Supplementary Table 8. Records of the number of sand flies collected in each sub-pool

Information about the number of female sand flies infected with sub-pools P1-P4 and masterpool at 2 and 9 days PBM for bar-seq analysis.
