## Supplementary figures and images for "Identification of transporters essential for survival of *Leishmania* promastigotes in the digestive tract of sand flies"

### Supplementary Figure 1.pdf

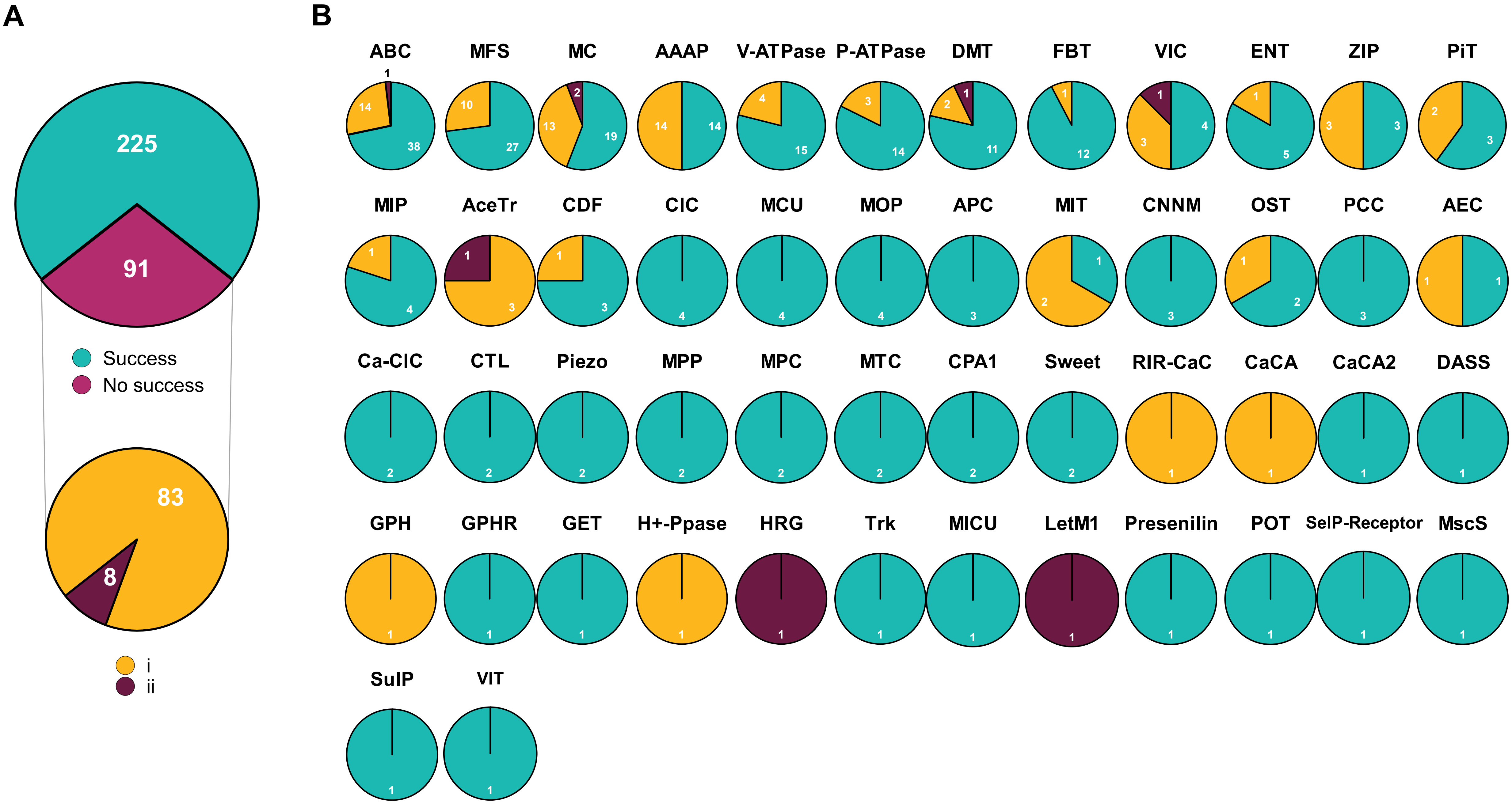


Supplementary Figure 1

### Supplementary Figure 3.pdf

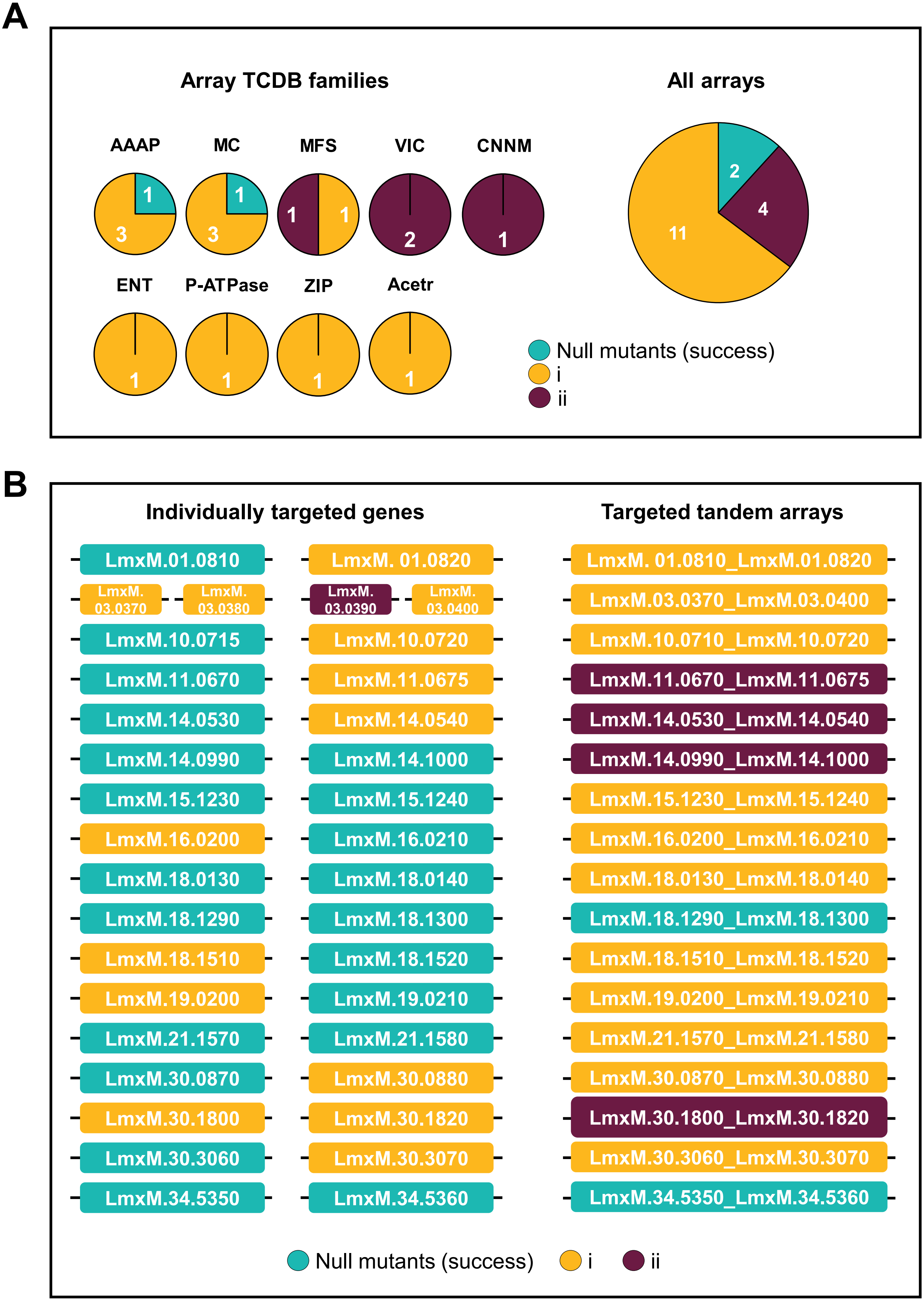


#### Supplementary Figure 3

### Supplementary Figure 4.pdf

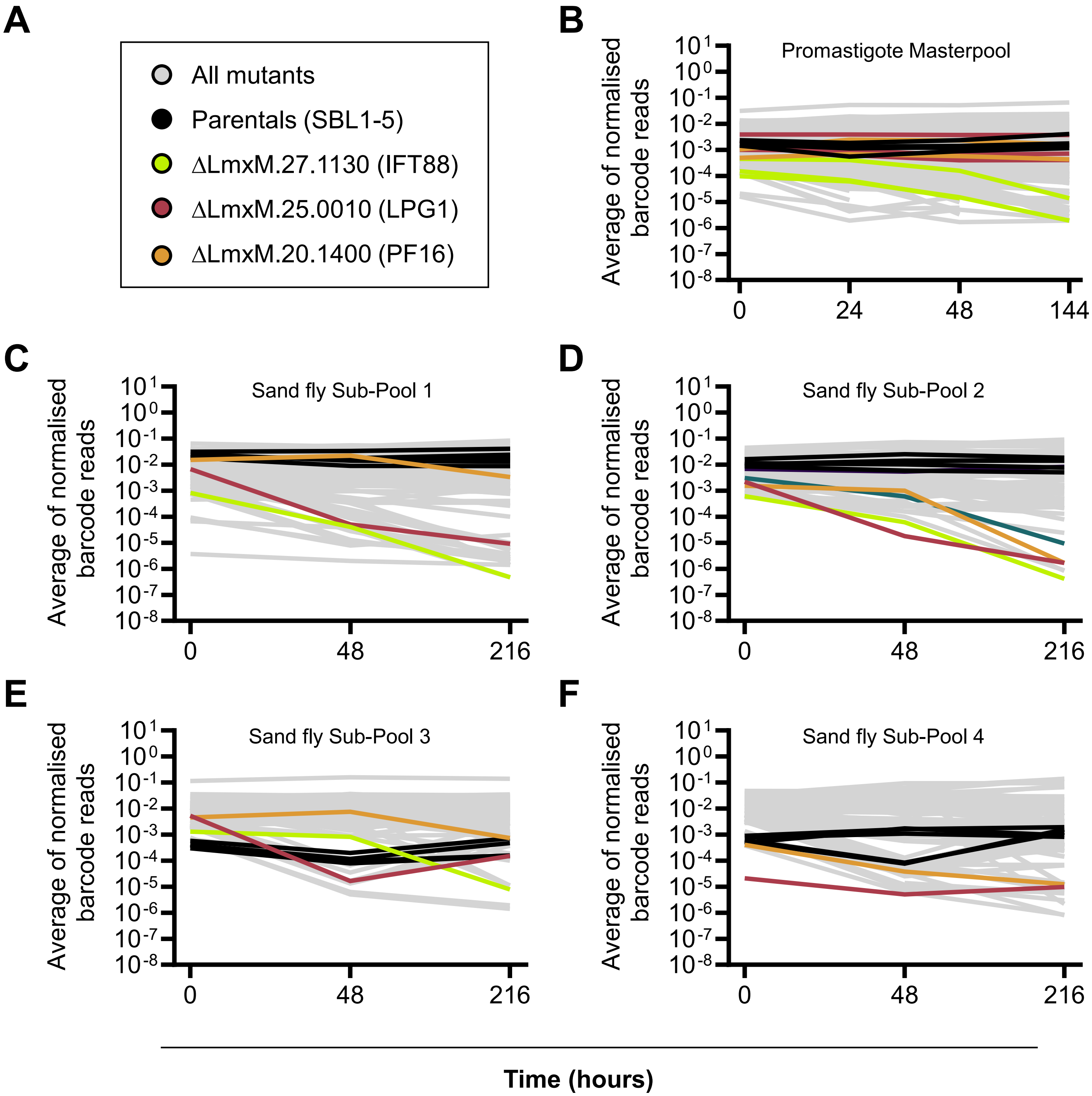


#### Supplementary Figure 4

### Supplementary Figure 5.pdf

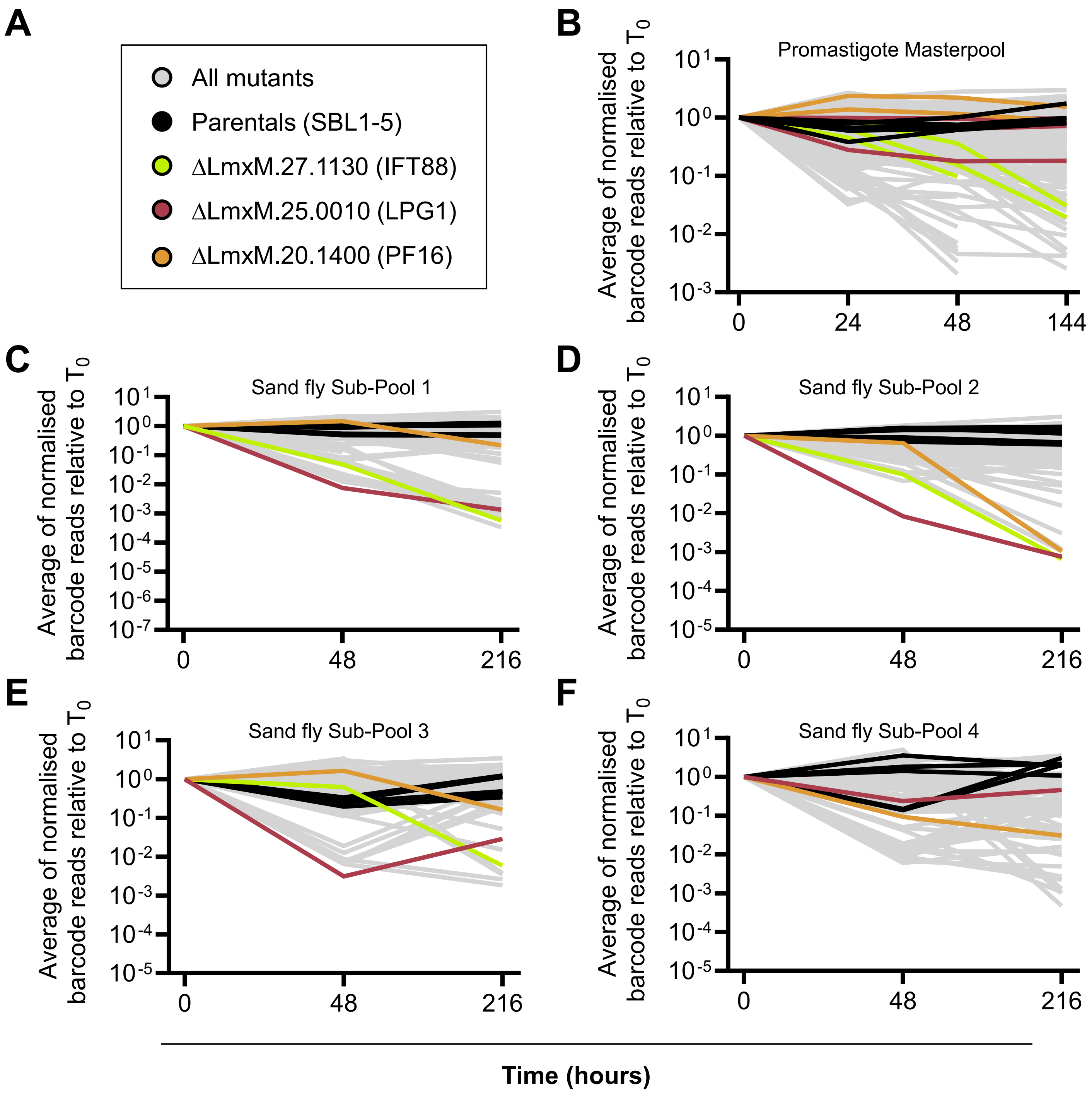


#### Supplementary Figure 5

### Supplementary Figure 6.pdf

A
B
C
D

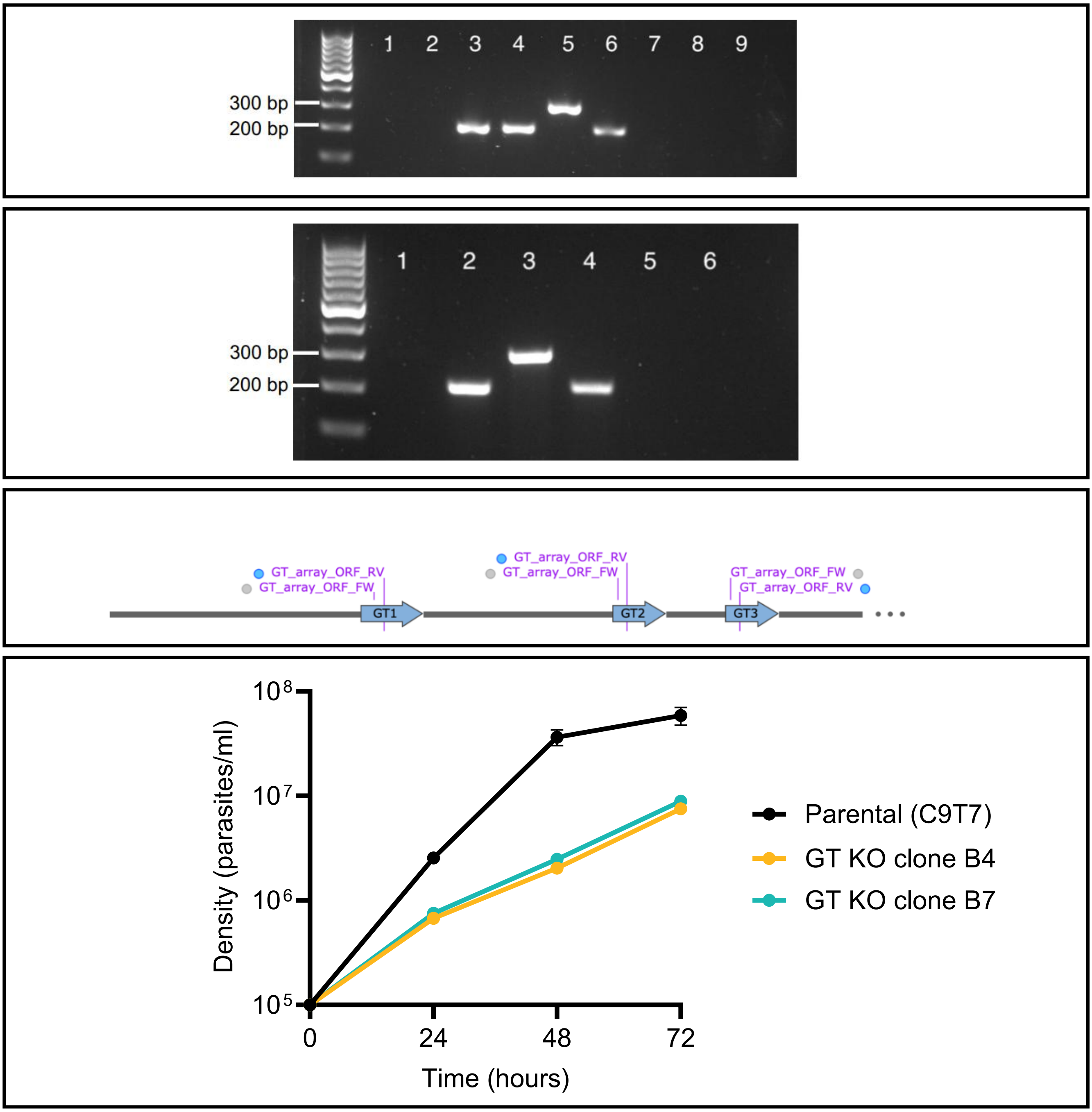


E

| Time period (hours) | Parental (C9T7) | GT KO Clone B4 | GT KO Clone B7 |
|---|---|---|---|
| 0 – 24 | 5.14 | 8.73 | 8.24 |
| 0 – 48 | 5.64 | 11.05 | 10.37 |
| 24 – 48 | 6.24 | 15.05 | 13.99 |
| 48 – 72 | 34.77 | 12.70 | 13.03 |

Supplementary Figure 6

### Supplementary Figure 7.pdf

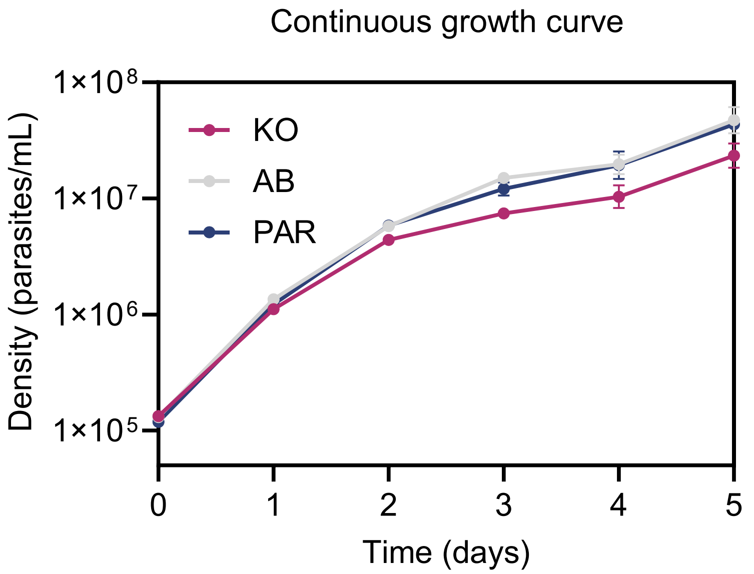


#### Supplementary Figure 7
